# Piezo1 as a force-through-membrane sensor in red blood cells

**DOI:** 10.1101/2022.08.10.503510

**Authors:** George Vaisey, Priyam Banerjee, Alison J. North, Christoph A. Haselwandter, Roderick Mackinnon

## Abstract

Piezo1 is the stretch activated Ca^2+^ channel in red blood cells that mediates homeostatic volume control. Here we study the organization of Piezo1 in red blood cells using a combination of super resolution microscopy techniques and electron microscopy. Piezo1 adopts a non-uniform distribution on the red blood cell surface, with a bias towards the biconcave “dimple”. Trajectories of diffusing Piezo1 molecules, which exhibit confined Brownian diffusion on short timescales and hopping on long timescales, also reflect a bias towards the dimple. This bias can be explained by “curvature coupling” between the intrinsic curvature of the Piezo dome and the curvature of the red blood cell membrane. Piezo1 does not form clusters with itself, nor does it co-localize with F-actin, Spectrin or the Gardos channel. Thus, Piezo1 exhibits the properties of a force-through-membrane sensor of curvature and lateral tension in the red blood cell.

## Introduction

Red blood cells (RBCs) experience significant mechanical forces in their lifetime as they traverse the mammalian circulatory system, squeezing through capillaries of smaller diameters than themselves (Chien 1987). Central to the ability of RBCs to deform in response to shear stress is their biconcave disk shape (Hodgkin and Lister 1827; Mohandas and Evans 1994). RBCs lack organelles and a transcytosolic network and their biconcave shape is instead imparted by the plasma membrane and the membrane skeleton, a 2D quasihexagonal network of actin filament (F-actin) nodes connected by (α1β1)2-spectrin tetramers bound to transmembrane proteins (Fowler 2013; Gratzer 1981). In response to shear stress a Ca^2+^ influx is activated in RBCs (Larsen et al. 1981; Dyrda et al. 2010). Ca^2+^ entry into RBCs modulates junctional protein interactions in the membrane skeleton (Nunomura 2006, 1) in addition to activating the Gardos channel (Gárdos 1958), a K^+^ channel, leading to K^+^ efflux and subsequent cell volume decrease (Danielczok et al. 2017), which together facilitate flexibility of the cell membrane.

Piezo1 was first suggested as the mechanosensitive Ca^2+^ channel in RBCs after identification of gain-of-function mutations in patients with dehydrated xerocytosis (Bae et al. 2013; Albuisson et al. 2013; Andolfo et al. 2013; Zarychanski et al. 2012). Subsequent studies using a hematopoietic PIEZO1 knockout mouse confirmed that the channel is responsible for stretch-induced calcium influx in RBCs (Cahalan et al. 2015). How Piezo1 is organized in the RBC membrane, however, is not known. The unique biconcave shape of the RBC poses interesting questions about how membrane curvature, which locally is known to be important for Piezo1 structure (Guo and MacKinnon 2017; Lin et al. 2019; Haselwandter and MacKinnon 2018), might globally impact Piezo1 distribution and function. It has been proposed that due to its intrinsic curvature, Piezo1 might tend to concentrate in the dimple region of RBCs (Svetina, Švelc Kebe, and Božič 2019), but direct experimental evidence is lacking. More broadly, how Piezo channels are organized in cell membranes is poorly understood. Based on electrophysiological recordings in overexpressing cells (Gottlieb, Bae, and Sachs 2012) or fluorescence microscopy of a stable cell line (Ridone et al. 2020), some authors have reported clustering of Piezo1, but channel densities in both cases are likely non-physiological. Extensive patch-clamp studies and simulation of Piezo1 densities in neuro2Acells, which express Piezo1 natively, led others to the conclusion that Piezo1 channels are homogenously expressed across cell membranes and function as independent mechanotransducers (Lewis and Grandl 2021).

Additionally, it is unclear what role the actin cytoskeleton plays in organizing and gating the Piezo1 channel in cells. Piezo1 can be pressure-activated in membrane blebs lacking cytoskeleton (Cox et al. 2016) and exhibits spontaneous openings in asymmetric lipid droplet bilayers (Syeda et al. 2016), consistent with a “force-from-lipids” model of activation (Martinac, Adler, and Kung 1990). Others claim that Piezo1 is tethered to the actin cytoskeleton and is gated by a “force-from-filament” mechanism (Wang et al. 2022).

Here we set out to image and analyze the organization of Piezo1 channels in RBC membranes. The small size of RBCs and their inherent autofluorescence makes them refractory to conventional fluorescence microscopy, so we employed more sensitive super resolution microscopic methods. Fluorescent Piezo1 spots can be resolved into single Piezo1 channels which, along with electron microscopy studies, confirm that they do not cluster in RBC membranes. Although not clustering, we find a non-uniform distribution of Piezo1 in the RBC membrane with enrichment at the dimple, which can be rationalized through the energetic coupling of the intrinsic curvature of Piezo1 to the curvature of the surrounding membrane. Single particle tracking (SPT) studies of Piezo1 demonstrate its lateral mobility in the plasma membrane, consistent with co-immunostaining analysis, suggesting that Piezo1 is not tethered to the membrane skeleton. Mobility of Piezo1 in the RBC membrane permits Piezo1 to explore the curvature landscape of the RBC surface and may allow Piezo1 to respond to local and global changes in membrane curvature as well as membrane tension.

## Results

### Expression and localization of Piezo1 in red blood cells

To detect Piezo1 in RBCS a PIEZO1-HA-knock-in mouse line was generated (Jackson Laboratories) in which an HA epitope tag was inserted into an extracellular loop of the PIEZO1 gene. The PIEZO1-HA-knock-in channel functions similarly to the WT channel in electrophysiological recordings of HEK cells and could be specifically detected by immunostaining (Supplementary Fig. 1). Importantly, only RBCs from the PIEZO1-HA-knock-in mice and not wild type mice showed fluorescent signal when immunostained with an anti-HA antibody (Fig. 1*A*), confirming clear expression of Piezo1 in RBCs. As shown in 3D reconstructions, single optical sections and volume renderings from reconstructed 3D structured illumination microscopy (3D-SIM) images (Gustafsson 2000; Heintzmann and Huser 2017) of immunostained RBCs, Piezo1 channels appear as spots (Fig. 1 *A*, *B* and Supplementary Fig. 2). These spots are distributed on the RBC membrane, here contoured by phalloidin staining of actin, which forms part of the actin-spectrin network bound to the cytosolic face of the RBC membrane (Ballas and Krasnow 1980) (Supplementary Fig. 2). 3D reconstructions from z-stacks reveal ∼80 Piezo1 puncta per RBC (Fig. 1*C*), suggesting a relatively low copy number of channels per RBC, similar to estimations of Gardos channel number per RBC (Lew, Muallem, and Seymour 1982; Brugnara, De Franceschi, and Alper 1993) and in contrast to very abundant RBC membrane proteins such as Band3, which is estimated to have a copy number of 1 million per cell, (Fairbanks, Steck, and Wallach 1971). Supportively, our patch-clamp recordings of RBCs occasionally identified single channel currents that are stretch-activated and have a conductance consistent with other reports of Piezo1 (Coste et al. 2010; del Mármol et al. 2018; Harraz et al. 2022) (Fig. 1*D*).

**Figure 1.**
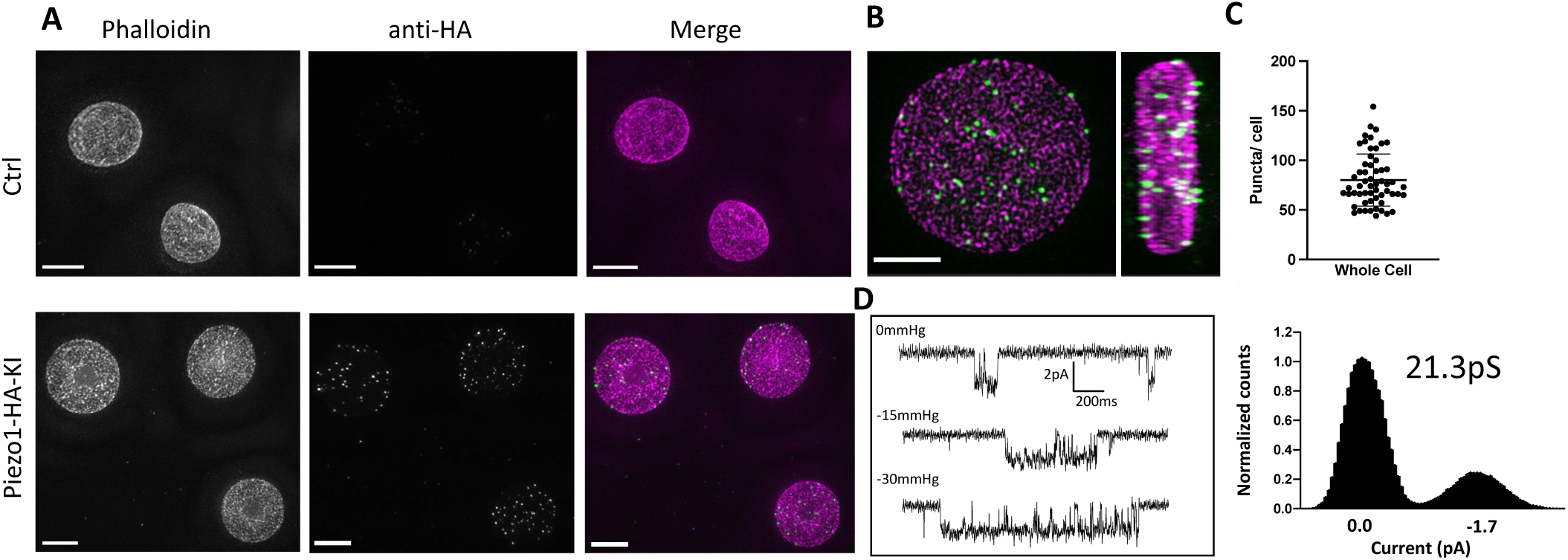
**A** Reconstructed 3D SIM images of RBCs, shown as maximum projections of Z-stacks, from control (Ctrl, top row) or Piezo1-HA-KI (bottom row) mice labelled using rhodamine phalloidin (magenta) and an antibody directed against the HA tag of Piezo1 (green). Scale bar, 3 μm. **B** 3D volume rendering of a representative labelled Piezo1-HA-KI mouse RBC displayed as XY (left) and XZ (right) axis views, showing the distribution of Piezo1. Scale bar, 2 μm **C** Quantification of the number of Piezo1 spots per cell from 3D SIM maximum projections. Mean ± SD = 80.2 ± 26 spots per cell. Numbers were calculated on individual cells (n=57) using Imaris spot detection. **D** Representative single channel traces from excised inside-out patch recordings of RBCs in symmetric 150mM NaMethanesulfonate. The patch was clamped at -80 mV and negative pressure (0 to -30mmHg) was applied to the patch pipette. Amplitude histogram (right) enables estimation of single channel conductance.

### Piezo1 does not cluster in RBC membranes

Whether Piezo1 channels cluster in membranes to form mechanosensory domains or are distributed more randomly as independent mechanosensors remains unknown. In 3D-SIM images of labelled intact RBCs, fluorescent Piezo1 spots are relatively uniform in size and do not appear to cluster (Fig. 1). However, our SIM data are diffraction-limited by an XY resolution of ∼120nm at best and thus each individual spot could feasibly contain a few Piezo1 channels (∼23nm diameter) tightly packed together. To address whether fluorescent spots represent single or multiple Piezo1 channels we therefore employed stimulated emission depletion (STED) microscopy (Hell and Wichmann 1994), which can achieve resolutions below 50nm and down to e.g. 22nm after deconvolution (Schoonderwoert et al. 2013), depending on the selected dyes.

Owing to the absorption of high-power lasers by heme (Schloetel et al. 2019), it was not possible to image intact RBCs by STED. Instead, RBCs adhered to poly-lysine coated coverslips were unroofed by a stream of isotonic buffer (Swihart et al. 2001), washing away hemoglobin and leaving patches of RBC membrane still bound to the actin-spectrin network (Fig. 2*A*). The approximate flat dimensionality of unroofed RBCs also enabled us to use facilitated image acquisition 2D STED mode to achieve maximal XY resolutions of ∼ 40 nm by full width half maximum (FWHM) in single images, closer to the size of a Piezo1 channel. Approximately 26 ± 7 (n= 12 cells) spots were observed per unroofed RBC, which equates to ∼ 0.6 puncta/μm^2^, consistent with the Piezo1 abundance observed in SIM images of intact RBCs. Piezo1 spots in 2D STED images of unroofed RBC membranes show a range of nearest neighbor distances (Fig. 2*B*) with an average nearest spot distance of 560 nm ± 130 nm (*n* = 12 cells). This is similar to the observed distribution of Piezo1 spots in 3D SIM images of intact cells (Fig. 2*C*), where the nearest spot distance is 540 nm ± 37 nm (*n =* 19 cells) and is consistent with fluorescent Piezo1 spots not clustering in the RBC membrane. Further cluster analysis of Piezo1 spots in 2D STED images by 2D spatial analysis (Andrey et al. 2010) shows no statistical difference between the observed spot distribution and a simulation of randomly distributed spots (Supplementary Fig. 3).

**Figure 2.**
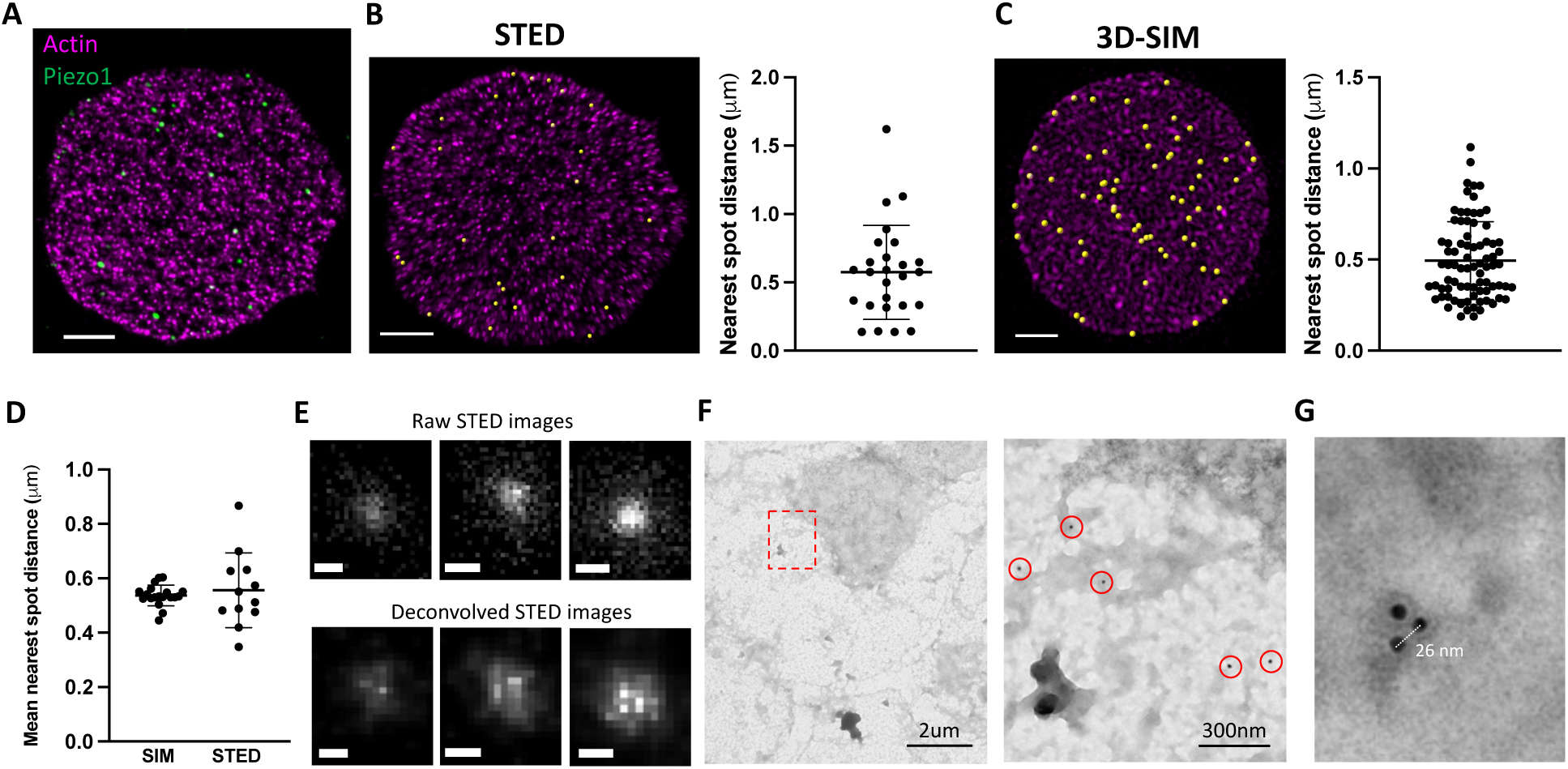
Piezo1 does not cluster in RBCs. **A** Representative 2D STED image of an unroofed RBC immunostained with an anti-HA tag antibody for Piezo1 (green) and labelled with phalloidin STAR 580 for F-actin (magenta). Scale bar = 1μm **B** 2D STED image of an unroofed RBC as in *A* where Piezo1 spots are detected by the spot detector function in Imaris and are colored yellow. The distance from each Piezo1 Piezo1 spot to the nearest Piezo1 in this cell was calculated in Imaris and is shown on the right. Scale bar, 1 μm. **C** The same analysis as in *B* was carried out on a 3D reconstruction from SIM imaging of immunostained RBCs. Scale bar, 1 μm. **D** The mean nearest spot distance was calculated for SIM and STED analyses and plotted. For SIM the mean is 540 nm ± 37 nm (n = 19 cells) and for STED the mean is 560 nm ± 130 nm (n = 12 cells). **E** Huygens deconvolution of Piezo1 puncta resolves into single, double and triple labeled spots where bright pixels are separated by ∼25 nm. Scale bar is 50 nm. **F** Negative-stain electron microscopy of an unroofed RBC immunostained with an anti-HA tag primary antibody and a secondary antibody conjugated to 18nm gold. Low magnification image (left) of an unroofed RBC (left) with a region highlighted in red imaged at medium magnification (right). 18nm gold particles corresponding to labelled Piezo1 channels are highlighted by red circles. As in fluorescence microscopy images, Piezo1 channels do not appear to cluster. **G** Negative stain electron microscopy image at high magnification of a triple-labeled Piezo1 channel.

Optimizing 2D STED image acquisition parameters, including imaging over multiple Z planes and employing adaptive illumination (Heine et al. 2017), as well as image processing through spherical aberration correction and iterative deconvolution (Huygens Professional) (Schoonderwoert et al. 2013) enabled an improvement of the lateral full width at half maximum (FWHM) resolution towards ∼ 25nm (Fig. 2*E*), approximating the diameter of the Piezo channel. Post-deconvolution, we observed that a significant number of fluorescent spots resolved into triplets of bright spots, consistent with labelling each arm of the trimeric structure of Piezo1 (Saotome et al. 2018; Guo and MacKinnon 2017) (Fig. 2*E* and Supplementary Fig. 4). In a single field of view we can often observe multiple triple-labeled Piezo1 channels (Supplementary Fig. 4). Single- and double-labeled Piezo1 channels were also observed. Thus fluorescent spots do not represent multiple Piezo1 channels clustered tightly together. The frequent occurrence of triple-labeled Piezo1 in our STED data suggests we are towards the upper limit of labelling. An important corollary to this conclusion is that, at least if the probability of observing a given number of antibodies bound to each Piezo1 channel is non-cooperative, we should expect only a small fraction of Piezo1 channels to be unlabeled, giving us confidence in our estimation of Piezo1 channel number per RBC (Fig. 1*C*). This settles inconsistencies in the available quantitative proteomics analyses of the RBC proteome (Bryk and Wiśniewski 2017; Ravenhill et al. 2019) and is in line with a study of highly purified erythrocytes, which estimates approximately one hundred Piezo1 channels per cell (Gautier et al. 2018). We further analyzed Piezo1 distribution in unroofed red blood cells by negative stain electron microscopy using an 18nm gold-conjugated secondary antibody (Fig. 2*F* and *G*). Gold particles were distributed over the unroofed RBC membrane without apparent clustering, comparable to the distribution of fluorescent spots in STED images. Gold particles were not observed in RBCs from control mice. By negative stain electron microscopy, we were also able to observe triplets of gold particles (Fig. 2*G*) with an inter-particle distance of ∼25nm, consistent with fully labelled trimeric Piezo1 channels. Thus, using multiple imaging approaches, we do not observe Piezo1 clustering in the RBC membrane.

### Piezo1 is enriched at the dimple region of RBC membranes

We were curious whether Piezo1 is evenly distributed throughout the RBC membrane or tends to localize to specific regions of the RBC biconcave disk. Because of its intrinsic curvature, it has been hypothesized that Piezo1 concentrates in the curved dimple of the RBC membrane (Svetina, Švelc Kebe, and Božič 2019). Other force-related proteins such as myosin IIA exhibit a non-uniform membrane density and are enriched at the dimple region (Alimohamadi et al. 2020). Using 2D slices from 3D-SIM images of intact RBCs we segmented the cells into the dimple and rim regions based on F-actin staining of the membrane (Fig. 3*A* and *B*) and quantified the number of Piezo1 puncta per μm^2^ surface area of each region using the cell imaging software Imaris (Fig. 3*C* and *D*). The dimple accounts for about 17% of the total RBC surface area. We find that the whole RBC and rim have similar Piezo1 puncta densities (0.50 ± 0.16 μm^-2^ and 0.46 ± 0.17 μm^-2^, respectively) while at the dimple region the density is higher (0.80 ± 0.31 μm^-2^). Thus, there is approximately a two-fold enrichment of Piezo1 at the dimple region of RBC membranes compared to the rim (Fig. 3*C*). This enrichment of Piezo1 at the dimple region of the RBC membrane is not due to enzymatic deglycosylation of RBCs, which we used to increase antibody binding to HA-tagged Piezo1 (Supplementary Fig. 5). To strengthen the hypothesis that it is the intrinsic curvature of Piezo1 that biases it towards the curved dimple region of the RBC we also analyzed the dimple vs. rim ratio of two other RBC membrane proteins that are not intrinsically curved: the Gardos channel (Lee and MacKinnon 2018) and Band3 (Vallese et al. 2022; Xia, Liu, and Zhou 2022; Arakawa et al. 2015). We find no statistically significant enrichment of the Gardos channel (KCNN4) or Band3 in the dimple of RBCs (Fig. 3*C*). Because we observe a range of relative enrichments of Piezo1 at the dimple we asked whether this correlates with the degree of RBC biconcavity, since we also observe a range of biconcavities via F-actin staining of the RBC membrane (Supplementary Fig. 5). The degree of biconcavity was parameterized as the ratio of the maximum to minimum heights of RBCs at a central xz slice (Fig. 3*D*). Indeed, with increasing RBC biconcavity, we observe a greater enrichment of Piezo1 at the dimple (Fig. 3*D*). If the distribution of Piezo1 in the RBC membrane is determined by the coupling of the channels’ intrinsic curvature to the global curvature of the membrane we would expect an unfavorable distribution of Piezo1 into membrane protrusions where the membrane curvature is opposing to that of Piezo1’s. Treatment of RBCs with NaSalicyate, which preferentially binds into the outer membrane leaflet, generates echinocytes (Li et al. 1999), RBCs that are characterized by “thorny” membrane protrusions. 3D-SIM imaging of labelled echinocytes shows exclusion of Piezo1 channels from such membrane protrusions, further supporting a mechanism of curvature coupling between the intrinsic curvature of Piezo1 and the surrounding membrane (Supplementary Fig.6).

**Figure 3.**
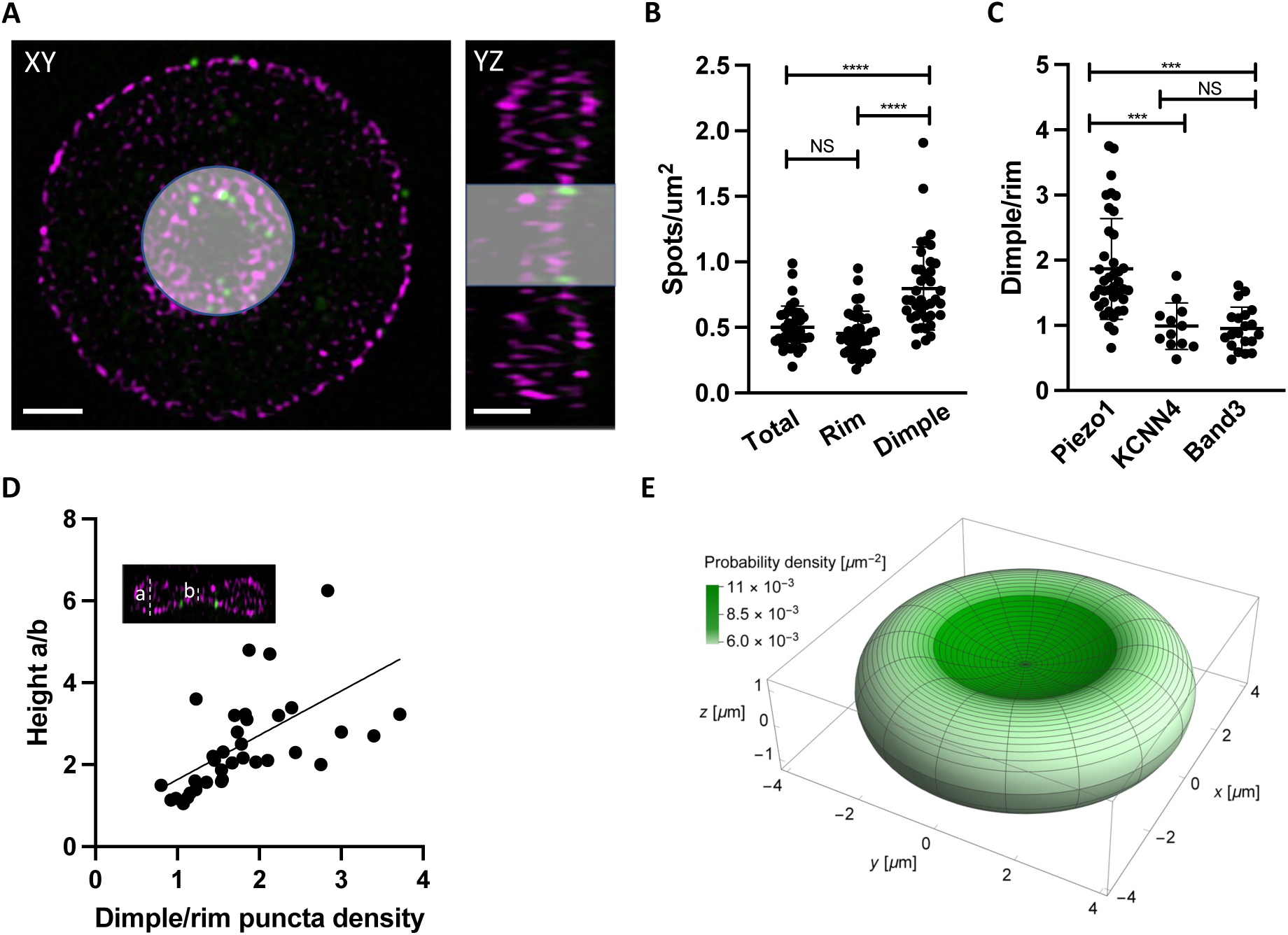
Piezo1 localization is enriched in the curved dimple region of the RBC. **A** Two dimensional slices in XY and YZ views from SIM images illustrating volume segmentation of RBCs and Piezo1 puncta distribution after labelling with an antibody against Piezo1-HA (green) and rhodamine-phalloidin for F-actin (magenta). Cells were segmented and masked in Imaris as shown. Scale bar, 1μm. **B** Distribution of Piezo1 puncta in RBCs. The RBC dimple has a higher density of Piezo1 puncta than whole RBCs (Total) (p=0.00019) and the rim region (p=0.00005) by the One-way Anova test with Tukeys Post Hoc analysis. N=39 cells **C** The ratio of Piezo1 spot density (1.87 ± 0.76, N= 39 cells) in the dimple and rim region of RBCs compared to that of KCNN4 (0.99 ± 0.36, N= 12 cells) and Band3 (0.95 ± 0.33, N=20 cells). The dimple over rim ratio is higher for Piezo1 than KCNN4 (p=0.0003) and Band3 (p=0.0005). **D** Scatterplot of ratio of Piezo1 puncta density in dimple over rim plotted against RBC biconcavity, measured as height ratio a/b (see inset). N=37 cells. Data are fit by linear regression with equation ! = 1.08’ + 0.55, R^2^ = 0.34 and P = 0.0001. **E** Probability per unit area for finding a Piezo1 channel at a given RBC membrane location, calculated from a physical model of Piezo1 curvature coupling and plotted over the RBC membrane surface. The RBC shape corresponds to the model of Beck for RBC discocytes (Beck 1978).

The relative abundance of Piezo1 in the dimple and positive correlation with the degree of biconcavity indicates that RBC membrane curvature might be the primary determinant of the Piezo1 distribution. The intrinsic curvature of Piezo1 suggests a possible mechanism, based on ‘curvature coupling’, for bias towards the dimple. To examine the plausibility of this mechanism, we developed a simple physical model of Piezo1 curvature coupling in which Piezo1 is represented as a spherical cap with area 450 nm^2^ and approximate radius of curvature 42 nm, the estimated asymptotic value in a planar membrane (PNAS, submitted). To describe the RBC shape, we used the model of Beck (Beck 1978) and calculated the mean curvature at each point on the surface (*Methods*). Then, using the Helfrich functional (Helfrich 1973), we calculated the minimum shape energy of the membrane surrounding Piezo1 in a vesicle that, if spherical, would have a radius of curvature equal to the reciprocal of the mean curvature at that point on the RBC surface. This procedure associates each point on the RBC surface with a membrane curvature energy which, when applied to the Boltzmann distribution equation, yields a probability distribution for the density of Piezo1 on the membrane surface. Further details of this calculation are outlined in *Methods*. The surface distribution of Piezo1 calculated according to this method is shown in Fig. 3*E*. The curvature coupling model predicts an approximately two-fold increase in the density of Piezo1 in the dimple relative to the rim, close to the experimentally observed ratio of densities. In the Beck model of RBC shape, the mean curvature at any point on the RBC surface ranges from 0.24 μm^-1^ (favorable to Piezo1 in the dimple) to about -0.52 μm^-1^ (unfavorable to Piezo1 in the rim). Because these mean curvatures for the RBC membrane are very small (compare these values to the approximate intrinsic curvature of Piezo1, 1/42 nm ≈24 μm^-1^), it was not immediately clear to us whether Piezo1 would be sensitive to surface curvature in the RBC. The correspondence between experiment and theory suggests that Piezo1 is indeed sensitive to RBC shape and adopts an equilibrium distribution by a mechanism of curvature coupling.

### Piezo1 is mobile within the red blood cell plasma membrane

The above distribution of Piezo1 according to the curvature coupling model suggests that Piezo1 ought to be laterally mobile so that it can explore the curvature landscape of the RBC membrane. However, the quasihexagonal actin-spectrin meshwork and large junctional protein complexes that make up the RBC membrane layer are known to corral and confine membrane protein diffusion (Sheetz 1983). Additionally, at least some membrane proteins, including roughly one third of Band3, are essentially immobilized by direct binding to actin-spectrin (Tomishige, Sako, and Kusumi 1998; Tsuji et al. 1988). We studied Piezo1 mobility in the RBC membrane with single particle tracking (SPT) using 40 nm gold nanoparticles conjugated to a secondary antibody under differential interference contrast (DIC) microscopy (Wu et al. 2019). The fast time resolution and ability to record over minutes without photobleaching has advantages for studying confined membrane protein diffusion (Tomishige, Sako, and Kusumi 1998; Daumas et al. 2003). Gold particles could be unambiguously identified on RBC membranes (Fig. 4*A* and Supplementary video 1). We imaged at 10Hz for 2 minutes. In recordings lasting 2 minutes we observed a range of Piezo1 diffusion trajectories (Fig. 4*B-D*).

**Figure 4.**
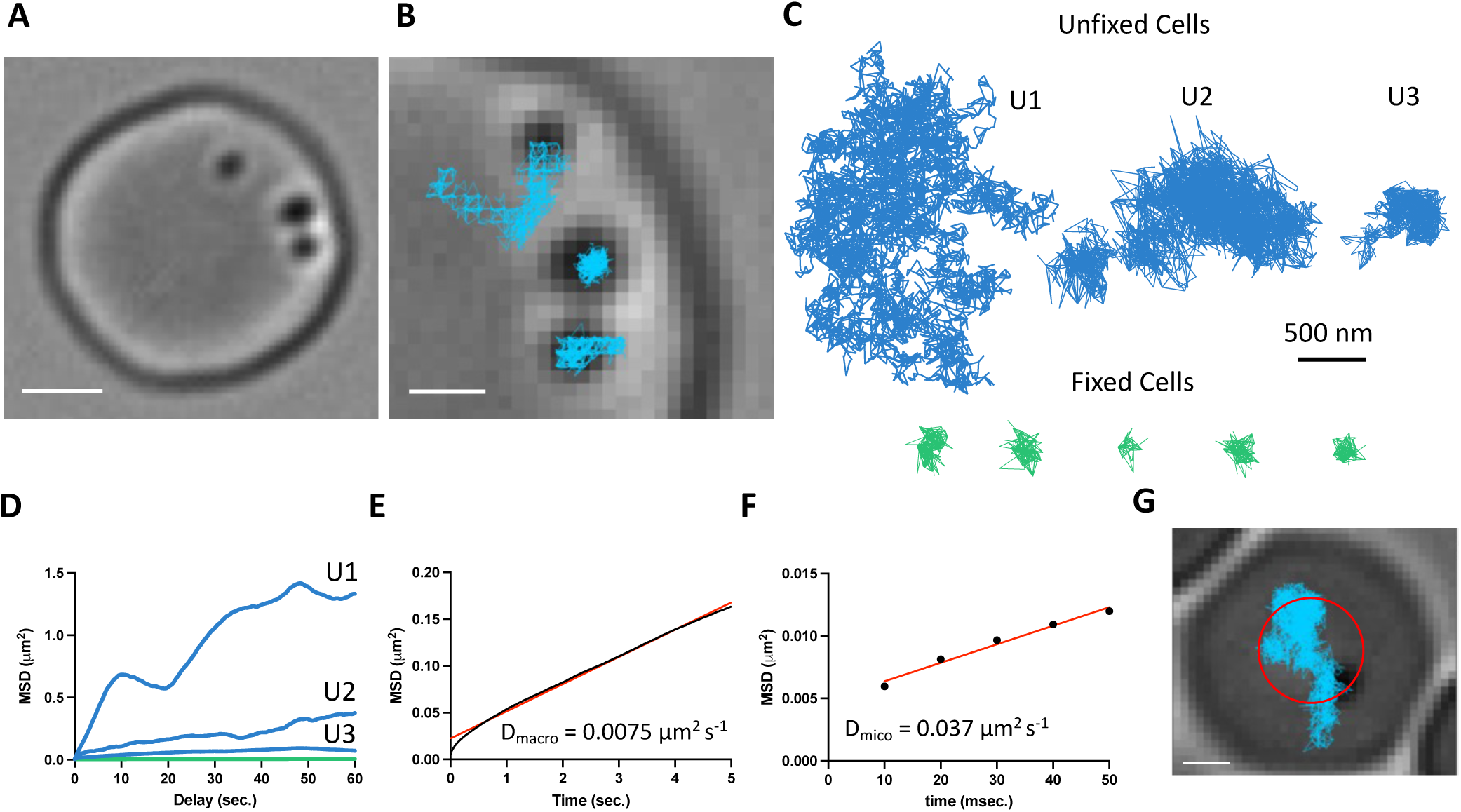
Gold-particle tracking of Piezo1 reveals its lateral mobility in the RBC membrane. **A** Example of a RBC imaged under DIC microscopy showing three 40 nm gold particle labelled Piezo1 channels. Scale bar, 2 μm. **B** Close up image of the gold particles with trajectories from a 2 minute recording at 100 Hz frequency. Scale bar, 1 μm. **C** Representative trajectories corresponding to distinct Piezo1 diffusive behaviors in unfixed conditions, shown in blue (trajectories U1, U2, and U3). The green trajectories correspond to labelled Piezo1 in cells fixed with PFA. **D** MSD against time for the first 60 seconds of the trajectories U1, U2, and U3 in *C* corresponding to unfixed conditions (blue) and an average of n = 5 fixed trajectories (green). **E** MSD against time for the first 5 seconds of an average of 14 trajectories. The data is fit to a straight line with a slope corresponding to a 2D diffusion coefficient of 0.0075 μm^2^ s^-1^. R^2^ for linear fit to data is 0.99. **F** MSD against time for the first 50 milliseconds of an average of 14 trajectories. The data is fit to a straight line with a slope corresponding to a 2D diffusion coefficient of 0.037 μm^2^ s^-1^. R^2^ for linear fit to data is 0.98. **G** Image of an RBC with a single gold labelled Piezo1 channel that was recorded for 1 hour at 1Hz frequency. Superimposed are the final tracking result (blue) and a circle (red) indicating the approximate segmentation of the RBC dimple. Scale bar, 1 μm.

Most trajectories exhibited confined diffusion, although sometimes Piezo1 appeared to be freely diffusing. Occasionally, we observed both types of diffusion in a single recording. In all cases, Piezo1 was more mobile than in control cells chemically fixed with paraformaldehyde, when analyzed by a mean squared displacement (MSD) analysis (Fig. 4*D*). For confined trajectories, we find that the confinement diameter is in the range of 200-600 nm. Although limited by the spatial resolution of our recordings, this range is largely consistent with the organization of the actin-spectrin meshwork we observe by 2D STED imaging of unroofed RBCs stained for actin or spectrin (Supplementary Fig. 7). In addition to smaller corrals of intact actin-spectrin, we find voids deficient in actin or spectrin staining, approximately 200-500 nm in size (Supplementary Fig. 6). Such voids in the actin-spectrin skeleton have also been observed by Stochastic Optical Reconstruction Microscopy (STORM) studies on RBCs (Rust, Bates, and Zhuang 2006). Plotting the average MSD of the first 5 seconds of Piezo1 tracks yielded an apparent macroscopic diffusion coefficient of 0.0075 μm^2^s^-1^ (Fig. 4*E*), which is similar to that of the mobile population of Band3 (Tomishige, Sako, and Kusumi 1998) and likely reflects occasional localized periods of confinement within actin-spectrin corrals over time. Microscopic diffusion was also analyzed by plotting the average MSD of the first 50 milliseconds of Piezo1 tracks before the channel will tend to become localized within confinement zones. This analysis yielded an apparent diffusion coefficient of 0.037 μm^2^s^-1^, consistent with measurements of freely diffusing Piezo1 in mouse neural stem progenitor cells (Ellefsen et al. 2019). The diffusion coefficient of a free ∼50nm gold nanoparticle in solution has been reported as ∼ 1 μm^2^s^-1^ (Giorgi et al. 2019), two orders of magnitude faster than the estimated microscopic diffusion coefficient of gold-labelled Piezo1 in our experiments. It is possible that a 40 nm gold particle may somewhat slow diffusion of membrane-embedded Piezo1, leading to an underestimation of its diffusion coefficient by our calculations. Nevertheless, Piezo1 appears laterally mobile within the RBC membrane.

Based on our imaging experiments of fixed RBCs that show an enrichment of Piezo1 in the dimple (Fig. 3), we tracked Piezo1’s preference for the dimple vs rim in real time by recording for an hour at 1Hz frequency (Fig. 4*G* and Supplementary video 2). We approximated the dimple as a circle with radius 1.13 μm, comparable to our segmentation based on F-actin staining (Fig. 3) and calculated the frequency at which gold-labelled Piezo1 is within 1.13 μm of an origin defined as the center of the RBC (Fig. 4*G*). We find that Piezo1 is observed within the dimple 81% of the time during this 1 hour recording, consistent with a preference of Piezo1 for the dimple over the rim.

### Piezo1 is not bound to the actin-spectrin meshwork

To further explore the behavior of Piezo1 in RBC membranes we analyzed its spatial proximity to the actin-spectrin meshwork by fluorescence microscopy. Proteins including ankyrin and protein 4.1 bind both to components of the meshwork and to proteins in the membrane, most commonly Band3 proteins, organizing the RBC membrane into functional microdomains composed of channels and transporters tethered to the actin-spectrin skeleton (Lux 2016).

Using co-immunostaining of unroofed RBC membranes, we asked whether Piezo1 is associated with these well-known complexes. Because of the small size of RBCs and the density of actin-spectrin at the membrane, super resolution microscopy was necessary to address this question. We employed 2D STED to image unroofed RBCs. To analyze spatial proximity of two fluorescent signals in our STED datasets we utilized optimal transport colocalization (OTC) (Tameling et al. 2021) analysis, which is more appropriate for STED microscopy than traditional pixel-based methods. First, we validated our experimental approach by co-immunostaining proteins of known complexes at the RBC membrane. There are three distinct known populations of Band3: one linked to spectrin via ankyrin, another bound to actin via protein 4.1 and other bridging proteins, and finally proteins that remain untethered to the actin-spectrin meshwork (Lux 2016; Kodippili et al. 2009). A significant fraction of Band3 fluorescent puncta are observed as overlapping or directly adjacent to both f-actin and spectrin puncta by STED microscopy, indicating close spatial proximity (Fig. 5*A*). In contrast, we do not observe a significant spatial proximity of Band3 and Piezo1, suggesting that if Piezo1 is tethered to actin-spectrin it is not as part of the Band3 complex. This is quantified by OTC analysis, where we plot the fraction of pixel intensities that is matched (colocalized) at distances less than the defined threshold distance. Following Nyquist sampling, the pixel size of 12.5 nm served as the lower limit of this threshold and the upper limit was set to 300nm (Fig. 5*B*). At a threshold of 100nm, for example, 0.6 or 60% of signals from Band3 complexes are considered proximal enough to specify as colocalized with both actin and spectrin, consistent with these the two aforementioned well-known populations of Band3 complexes. By contrast, only 0.15 or 15% of Piezo1 signals are considered proximal enough to specify as colocalized with Band3. We also co-immunostained Piezo1 with f-actin or spectrin to ask whether it is tethered to the RBC membrane skeleton. We do not observe significant spatial proximity between Piezo1 to either f-actin or spectrin by STED microscopy (Fig. 5*A*). At a threshold of 100nm we observe an OTC score of only 0.1 for Piezo (Fig. 5*B*). Thus, our data show that unlike Band3, Piezo does not tend to exhibit close spatial proximity to actin or spectrin and is unlikely to bind to the membrane skeleton, consistent with its observed mobility in tracking experiments.

**Figure 5.**
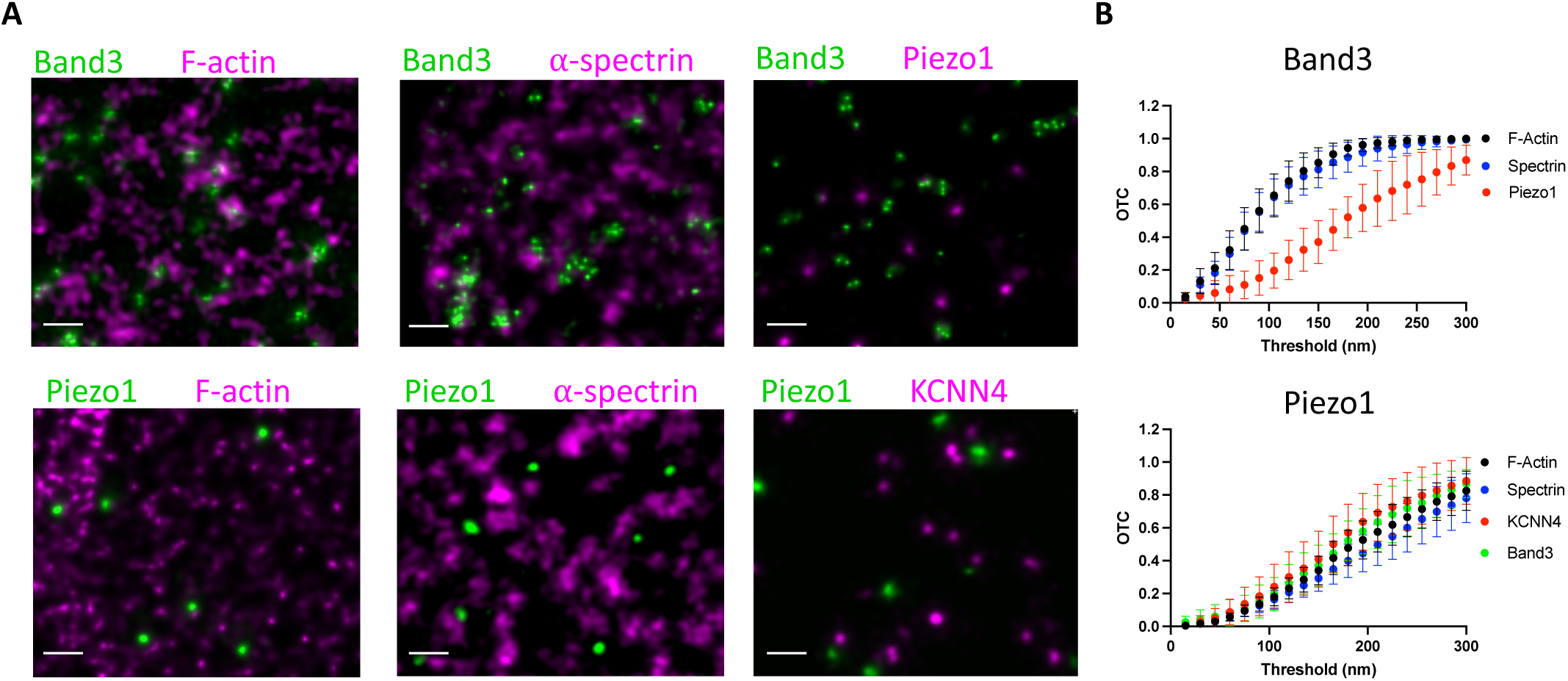
Piezo1 does not appear to interact with the membrane skeleton. **A** Exemplary STED microscopy images assessing interactions of Band3 (upper panels) and Piezo1 (lower panels) with known members of the RBC membrane skeleton as well with each other and with KCNN4. Close spatial proximity for Band3 with actin and spectrin can be observed in STED images but not for Piezo1. Scale bar, 200nm. **B** OTC analysis including 95% confidence error bars of STED microscopy image sections (Band3/F-actin, n=11; Band3/Spectrin, n=10; Band3/Piezo1, n=7; Piezo1/F-actin, n=10; Piezo1/Spectrin, n=7; Piezo1/KCNN4 n=8).

Based on a proposed mechanism of Piezo1-mediated Ca^2+^ influx activating the Gardos channel (Cahalan et al. 2015) (KCNN4), we also analyzed spatial proximity of these two proteins by STED. Piezo1 and KCNN4 do not appear to form complexes in RBCs (Fig. 5*A* and *B*), but it should be noted, given the small size of the RBC, that co-localization of Piezo1 and KCNN4 is not necessary for a functional coupling of the ion fluxes associated with these two channels (see *Discussion*).

## Discussion

Since its discovery (Coste et al. 2010), Piezo1 has been identified as a primary mechanosensitive molecule in a wide range of force-sensing physiological processes from cell and organ development (Sun et al. 2019; J. Li et al. 2014; Ranade et al. 2014) to blood pressure regulation (Zeng et al. 2018) and stem cell lineage choice (Pathak et al. 2014). How Piezo1 is organized in different cell types and how this might pertain to the sensing of different types of physiological force is currently not well understood. Here we studied Piezo1 organization in RBCs, both because of the clear importance of Piezo1 in RBC physiology (Bae et al. 2013; Zarychanski et al. 2012; Cahalan et al. 2015) and because of the unique cellular structure of RBCs, particularly their biconcave disk shape.

At the smallest level of Piezo1 organization, that is with itself, we find that Piezo1 does not cluster in the RBC membrane. By 3D SIM microscopy (Fig. 1) we observe diffraction limited fluorescent spots that could technically represent multiple, closely located, Piezo1 channels. To distinguish these spots as individual channels, we needed to employ 2D STED microscopy (Fig.2). As the power of the depletion laser was increased to improve resolution, the size of each spot decreased correspondingly, but never resolved into multiple spots, even when the achieved resolution was near the size of the Piezo1 channel. Thus, we conclude that each fluorescent spot, whether viewed by 3D-SIM or 2D STED, corresponds to a single Piezo1 channel. The average distance between Piezo1 channels is approximately 550 nm, although occasionally we observe 2-3 channels within 100 nm distances. This is in good agreement with studies of neuro2A cells, which natively express Piezo1, that also found that Piezo1 channels do not tend to cluster (Lewis and Grandl 2021). While clustering could affect Piezo1 gating through Piezo’s membrane footprint, the precise coupling of Piezo1 gating to clustering would depend on the specific scenario considered, and clustering does not necessarily facilitate gating (Haselwandter and MacKinnon 2018). A relatively homogenous membrane distribution of Piezo1 might be well suited to RBCs that are tumbling through solution, experiencing shear forces from all angles, and may contribute to the remarkable plasticity of RBC shapes under stress (Dupire, Socol, and Viallat 2012). Of course, our data on RBCs do not rule out the possibility of Piezo1 clustering in other cell types.

Because of the intrinsic curvature of Piezo1 we were curious about its distribution as a function of the curvature of the RBC membrane. Experimentally we were able to ask this question through 3D SIM microscopy, which yielded sufficiently high-resolution spatial information on Piezo1 channels while enabling easy assignment of the RBC dimple and rim membrane regions (Fig. 3). Per μm^2^ area of membrane we find an almost 2-fold enrichment of Piezo1 in the dimple relative to the rim. Our experimental data can be rationalized by a physical model of curvature coupling between the intrinsic curvature of the Piezo1 dome and the surrounding membrane curvature of the RBC along its biconcave surface (see *Methods*). Whilst the curved biconcave shape of RBCs is unique, the good agreement between our experimental data and theoretical prediction based on energetic curvature coupling would suggest an inherent relationship between Piezo’s intrinsic curvature and the surrounding membrane that is broadly relevant to other cell types. This is because the curvature coupling theory, which was developed with experiments on lipid bilayer vesicles, depends only on the bending elasticity of membranes and the intrinsic curvature of Piezo1, not on the unique shape of RBCs. It will be interesting to explore whether Piezo1, for example, is enriched in membrane invaginations or other curvature-matched membrane regions in different cell types. Curvature-sensing of membrane proteins by curvature matching has been well documented for other intrinsically curved proteins such as Bin/Amphiphysin/Rvs (BAR) (Simunovic et al. 2015; Tsai et al. 2021) proteins. More recently, it has been shown that glycosylation can determine the membrane curvature sensing properties of some proteins (Lu et al. 2022). It is unclear from our current data whether the enrichment of Piezo1 in the RBC dimple is simply a corollary of the curved structure of Piezo1, which determines the channel’s tension sensitivity as described by the membrane dome model (Guo and MacKinnon 2017; Haselwandter and MacKinnon 2018), or whether it also has some important consequences for physiological force sensation of RBCs. In this context it is noteworthy that the force-generating protein myosin II A also shows an enrichment in the RBC dimple relative to the RBC rim (Alimohamadi et al. 2020).

Consistent with its ability to equilibrate along the RBC surface as a function of membrane curvature, we find that Piezo1 exhibits lateral mobility within the RBC plasma membrane. We used gold nanoparticle tracking under DIC (Fig. 4) so we could record for minutes to an hour without being limited by photobleaching and at fast time resolution (100Hz). This enabled us to record both the confined (<500 nm) diffusion of Piezo1 as well as Piezo1’s “hopping” into more freely diffusing behavior. The spatial scales associated with the confined diffusion regime most likely reflect the compartment sizes of the actin-spectrin meshwork we observe by 2D STED. Such a model of confined diffusion set by actin-spectrin is well documented in RBCs (Tsuji et al. 1988) and other cell types (Fujiwara et al. 2016). Similar to observations by others using STORM microscopy (Pan et al. 2018), we observe ∼ 200-500 nm voids that are deficient in actin or spectrin signal by 2D STED. Similar nanoscale “defects” in the actin-spectrin meshwork have also been observed by atomic force microscopy imaging of RBCs (Nowakowski, Luckham, and Winlove 2001) and may serve as weak points in the meshwork that facilitate fast RBC remodeling during shear stress. Such defects may also permit faster diffusion of transmembrane proteins such as Piezo1.

Our double labelling experiments suggest that Piezo1 does not form a complex with actin or spectrin, which is consistent with Piezo1’s confined diffusion within the actin-spectrin meshwork compartments, and should be contrasted with the near-immobilization observed for proteins such as CFTR and Band3 (Kodippili et al. 2009, 3; Haggie et al. 2006), which both form complexes with actin. Additionally, our data are not consistent with Piezo1 being tethered to the underlying actin and activated by a “force-from-filament” mechanism as has been traditionally described (Cox, Bavi, and Martinac 2019; Zhang et al. 2015). Of course, the actin-spectrin meshwork organizes the RBC membrane into compartments whose boundary conditions could impact how changes in local curvature and lateral tension of the membrane gate Piezo1. This could also explain how chemical agents that disrupt the actin cytoskeleton affect Piezo1 currents in other cell types (Gottlieb, Bae, and Sachs 2012; Wang et al. 2022), that is through changes in organization of the membrane rather than disruption of a direct tether to the channel itself.

Additionally, we asked whether Piezo1 might be co-localized with the Ca^2+^-activated K^+^ (Gardos) channel. Ca^2+^ flux into the RBC through Piezo1 and subsequent activation of the Gardos channel is thought to be the initial step in mechanically activated RBC volume regulation (Danielczok et al. 2017; Cahalan et al. 2015). We do not, however, observe significant spatial proximity between Piezo1 and Gardos channels. Given the small size of RBCs, approximately 1 μm thick in the dimple and 2 - 3 μm thick in the rim, and the Piezo1 density of around 0.5 per μm^2^, it seems that co-localization is probably not necessary for rapid opening of the Gardos channel following Piezo activation. For diffusion in 3 dimensions, the mean dispersion time can be approximated by " = ^!!^, % the mean distance and & the diffusion coefficient. The apparent diffusion coefficient for intracellular Ca^2+^ is in the range 13 – 65 μm^2^*/s* (Nakatani, Chen, and Koutalos 2002), meaning that Ca^2+^ should equilibrate within tens of ms within a RBC. In larger cell types, co-localization of Piezo1 with other channels might be necessary to elicit a functional coupling of ion fluxes. In rat ventricular myocytes, for example, Piezo1 has recently been shown, by diffraction-limited confocal microscopy, to co-localize with transient receptor melastatin (TRPM4) channels (Yu et al. 2022).

The experimentally determined values of the intrinsic curvature and the bending stiffness of Piezo1 in lipid bilayers suggest that the curvature differences across the unperturbed RBC membrane are, alone, insufficient to substantially change the shape of Piezo1, and thus are not expected to open its pore (PNAS, submitted). However, that Piezo1 distributes on the surface of RBCs by sensing the membrane curvature, by corollary, means that it must have the capacity to sense lateral membrane tension by a force-through-membrane mechanism. To see this, consider that Piezo1 has been shown to flex, i.e., change its curvature, as a function of local membrane curvature (PNAS, submitted). Consider also that membrane tension changes membrane curvature by flattening it. In this manner, curvature sensing and lateral membrane tension sensing are inextricably linked. This linkage lies at the heart of the membrane dome model of Piezo gating (Guo and MacKinnon 2017; Haselwandter and MacKinnon 2018; PNAS, submitted). When RBCs squeeze through small capillaries, the membrane shape as well as the transmembrane pressure change transiently. The resulting changes in membrane curvature and lateral membrane tension can both, according to the membrane dome model of Piezo gating, open Piezo1’s pore to trigger Ca^2+^ entry.

## Methods

### HA-tag knock in mice

C57/BL6 background mice heterozygous for an HA tag knocked into the coding sequence after amino acid position 893 of PIEZO1, generated by CRISPR/Cas9 technology, were purchased from Jackson Laboratories. Heterozygous mice were bred to generate mice homozygous for the HA tag knock in and were genotyped by PCR amplification around the 893 insertion site and gel electrophoresis analysis of the amplified band size.

### Preparation of blood

Adult mice (>3 months old) were anesthetized by isoflurane inhalation and retro-orbital bleeds were taken using a heparinized micro-hematrocrit capillary tube (Fisherbrand). Whole blood was washed in DPBS three times by centrifugation at 1000 x *g* to separate RBCs from the buffy coat.

### Electrophysiological recordings

Washed RBCs were diluted 1:1000 and plated on uncoated petri dishes containing bath solution consisting of 140 mM NaMethanesulfonate, 10 Hepes pH 7.4. RBCs were imaged with a Nikon eclipse DIC microscope at 40X magnification. Pipettes of borosilicate glass (Sutter Instruments; BF150-86-10) were pulled to ∼15-20 MΩ resistance with a micropipette puller (Sutter Instruments; P-97) and polished with a microforge (Narishige; MF-83). The pipette was filled with identical bath solution. Recordings were obtained with an Axopatch 200B amplifier (Molecular Devices), filtered at 1kHz and digitized at 10 kHz (Digidata 1440A; Molecular devices). Gigaseals were obtained by applying a suction pulse (-10mmHg) using a high-speed pressure clamp (ALA scientific). Holding at negative voltages often facilitated seal formation. To obtain excised inside-out patches, attached cells were lifted briefly to the air-water interface which about 50% of the time would remove the cell but leave an intact inside-out membrane patch in the pipette.

### Immunofluorescence of intact RBCs using 3D-structured Illumination microscopy (SIM)

Washed RBCS were deglycosylated by incubation with 10% PNGase F (New England Biolabs) at 37°C for 2 hours, which we found increased antibody binding. RBCs were then diluted 1:50 in DPBS with 4% PFA and incubated overnight at room temperature to fix. Fixed cells were washed three times in DPBS by centrifugation at 1000 x *g* for 5 mins and then permeabilized in DPBS + 0.3 % Triton X-100 for 15 mins. Permeabilized cells were then blocked in DPBS + 4% BSA + 1% normal goat serum (blocking buffer) overnight at 4°C. Permeabilized and blocked RBCs were then incubated with a rabbit anti-HA primary antibody (3724; Cell Signaling Technology) diluted in blocking buffer for 1 h at room temperature, washed three times in blocking buffer and then incubated in 1:500 dilution of Alexa-488-conjugated goat anti-rabbit secondary (ab150077; Abcam) mixed with 1X rhodamine-phalloidin (Invitrogen) for 1h, followed by washing three times in blocking buffer. For 3D-SIM imaging of KCNN4 and Band3 in RBCs, 1:100 dilution of rabbit anti-Band3 primary (Proteintech) and 1:500 dilution of rabbit anti-KCN44 (76647, Invitrogen) were used. 1:500 dilution of Alexa-488 conjugated goat anti-rabbit (ab150077; Abcam) was used as the secondary along with 1X rhodamine-phalloidin (Invitrogen).

3D-SIM images were acquired using a DeltaVision OMX V4/Blaze system (Cytiva) fitted with an Olympus 100x/1.40 NA UPLSAPO oil objective and Photometrics Evolve EMCCD cameras. 488 and 568 nm laser lines were used for excitation and the corresponding emission filters sets were 528/48 and 609/37 nm respectively. Image stacks were acquired with an optical section spacing of 125 nm. SI reconstruction and Image Registration were performed with softWoRx v 6.1 software using Optical Transfer Functions (OTFs) generated from Point Spread Functions (PSFs) acquired from 100 nm green and red FluoSpheres and alignment parameters refined using 100 nm TetraSpeck beads (Invitrogen).

Imaris version 9.9 was used for analyzing 3D-SIM images. Detection of Piezo1 fluorescent spots was performed with Imaris spot detection with an estimated XY diameter of 120nm, based on measurements of fluorescent spots in 2D slices and the approximate XY resolution limit of 3D- SIM experiments. Spots were mostly uniform in size with a maximum XY diameter of 180nm.

The intensity threshold for spot detection was set to between 3000-5000 based on excluding detection of weak background fluorescent signal on the coverslip surface.

### Echinocyte formation

Washed RBCs that were deglycsoylated were incubated in DPBS + 10mM NaSalicyate for 30mins at 37°C to generate echinocytes before fixation and immunostaining as described above.

### Immunofluorescence of unroofed RBCs using Stimulated Emission Depletion microscopy (STED)

Washed RBCs that were deglycosylated as above were plated onto poly-D-lysine coated high performance coverslips and allowed to adhere at room temperature for 15 min. Plated RBCs were unroofed by applying a stream of 10ml DPBS through a 20 gauge needle at a ∼20° angle (Swihart et al. 2001). Unroofed RBCs were fixed in DPBS with 4% PFA for 15 min and then washed three times in DPBS before blocking in blocking buffer for 30 mins. This was followed by immunostaining with 1:500 dilution of rabbit anti-HA primary (3724; Cell Signaling Technology) for 1 h at room temperature, washing three times in blocking buffer and then incubating in 1:500 dilution of anti-rabbit STAR RED secondary (Abberior) mixed with 1X phalloidin STAR 580 (Abberior) for 1h. Immunostained unroofed cells were then, again, washed three times in blocking buffer and mounted in uncured Prolong Diamond Antifade mountant (Invitrogen) before imaging. For co-immunostaining experiments, additional antibodies used were as follows: 1:100 dilution of rabbit anti-Band3 primary (Proteintech), 1:100 dilution of mouse anti-alpha 1 Spectrin (ab11751) and 1:500 dilution of rabbit anti-KCNN4 (76647, Invitrogen). For co-immunostaining experiments with rabbit primary antibodies, rat anti-HA (Roche) primary was used to label Piezo1 and in all cases the species-appropriate STAR RED and STAR 580 (Abberior) secondaries were used.

STED microscopy was performed using a Facility Line STED microscope (Abberior Instruments) equipped with Olympus IX83 stand, Olympus UPLXAPO 100x/1.45 NA oil objective, pulsed excitation lasers with time gating (405, 488, 561, and 640 nm) and a 775 nm pulsed STED depletion laser, 4 Avalance Photo Diode detectors, adaptive illumination packages (DyMIN/ RESCue), and a deformable mirror for correction of spherical aberrations. Abberior Imspector software version 16.3.14287-w2129 with Lightbox interface was used for image acquisition.

Fluorophore excitation was facilitated at 580 nm (Phalloidin STAR 580) and 640 nm (STAR RED), whereas depletion was achieved using a wavelength of 775 nm. A pixel size of 12.5 nm was used in 2D STED mode. Excitation laser power, depletion laser power, line averaging/ accumulation, and pixel dwell time were optimized for balancing signal-to-noise ratio (SNR), and resolution, while minimizing photobleaching. Adaptive illumination techniques RESCue (Staudt et al. 2011) and DyMIN (Heine et al. 2017) were employed to reduce photobleaching and increase SNR.

Acquired STED images were saved as OBF files, which were deconvolved using theoretical point spread function (PSF) modeled from microscopy parameters in Huygens Professional version 22.04 (Scientific Volume Imaging). Imported images were cropped as needed, microscopic parameters were edited for refractive index mismatch correction, the excitation fill factor was set to 1.38 per software developer recommendations, and the STED saturation factor was tested within a range of 40-80, as per vendor recommendations, for optimal resolution. The deconvolution wizard was used to estimate background manually in the raw images and the acuity option was set for optimal image sharpness. Deconvolved images were exported as 16- bit TIFF files.

Cluster analysis on 2D STED images was performed using the spatial statistics 2D/3D ImageJ plugin (Andrey et al. 2010).

### Image analysis in Imaris

Imaris version 9.9 was used for analyzing fluorescent images. Detection of Piezo1 fluorescent spots was performed with Imaris spot detection with an estimated XY diameter of 120nm, based on measurement of fluorescent spots in 2D slices and the approximate XY resolution limit of 3D-SIM experiments.

### Negative stain electron microscopy of unroofed RBCs

Washed and deglycosylated RBCs were diluted 1:100 and added to carbon coated 400 mesh copper grids (CF400-CU, Electron Microscopy Sciences) that had been glow discharged and pre-coated in poly-d-lysine solution. After 30 mins incubation, RBCs were unroofed in DPBS as described above. Unroofed RBCs were blocked for 30 mins at room temperature in blocking buffer and then incubated with 1:200 rabbit anti-HA primary antibody (3724; Cell Signaling Technology) for 1 h at room temperature. Unroofed cells were then washed three times in blocking buffer and incubated with 1:20 diluted 18nm gold-conjugated goat anti-rabbit secondary antibody (Jackson Laboratories) for 1 h at room temperature. After rinsing three times in blocking buffer, grids were then washed three times in 150NaCl, 20 Hepes 7.5 to remove phosphate before staining in 1% uranyl acetate for 1 min. Images were collected on a Tecnai G2 Spirit BioTWIN Transmission Electron Microscope.

### Probability distribution calculation of Piezo1 along the RBC membrane

The Beck RBC model was used to describe the surface of a biconcave disk with diameter 8.0 μm, dimple thickness 1.0 μm, maximum rim thickness 2.5 μm, and area 138 μm^2^ (Beck 1978). The mean curvature at each point on this surface was calculated as 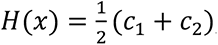, where *c*_1_ and *c*_2_ are the principal curvatures, which are functions of the position *x* on the surface (Carmo 2018). The following calculation was carried out to assign a (relative) membrane energy, and associated probability density, to Piezo1 at each position *x* on the RBC surface. The aim of this calculation is not to provide a detailed description of RBC membrane shape but, rather, to test whether Piezo1 localization can couple to curvatures on the RBC surface that are 1-2 orders of magnitude smaller than the Piezo dome curvature. In particular, Piezo1, modelled as a spherical cap (dome) with area 450 nm^2^ and radius of curvature 42 nm (PNAS, submitted), was placed into a vesicle whose radius of curvature, if it were spherical, equals 1/*H*(*x*). We note that the above conditions specify the area of the vesicle, which equals the free membrane area plus 450 nm^2^. For positions on the RBC surface at which the orientation (sign) of the RBC curvature matches the orientation of the Piezo1 intrinsic curvature (concave side facing outside the cell) Piezo1 was placed into the vesicle with an inside-out orientation, i.e., matching the orientation of the vesicle curvature. Otherwise, Piezo1 was placed into the vesicle with an outside-out orientation. The energy of the vesicle free membrane (outside the perimeter of the Piezo dome) was calculated through minimization of the Helfrich membrane bending energy

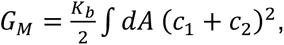

where the lipid bilayer bending modulus *K_b_* ≈ 20 *k*_B_*T* and the integral is carried out over the entire vesicle free membrane, subject to the boundary conditions at the Piezo dome-free membrane interface mandated by the shape of the Piezo dome (Helfrich 1973; PNAS, submitted). The resulting energy as a function of RBC curvature (and, thus, as a function of the position *x* on the RBC surface) was applied to the Boltzmann distribution equation to calculate the probability per unit area of observing Piezo1 across the RBC surface. This probability density is graphed on the surface of the Beck RBC model in Fig. 3*E*. Three major assumptions underlie our calculation of the curvature coupling energy. First, Piezo1 is treated as a spherical cap.

Second, for each position on the RBC surface the two principal curvatures, *c*_1_ and *c*_2_, are used to define a radius of curvature equal to the reciprocal of the mean curvature. Both the first and second assumption simplify the problem by imposing radial symmetry on the system of the Piezo dome and its surrounding membrane. The third assumption is that Piezo1 is sensing the curvature associated with the overall shape of the RBC, rather than the curvature of a locally specified surface (small membrane compartment) with nearby fixed boundaries.

We also carried out a joint energy minimization of the free membrane as described above while simultaneously permitting Piezo1 to flex, i.e., to change its radius of curvature, as part of the energy minimization. This calculation was made possible through our recently determined bending modulus of the Piezo dome itself (PNAS, submitted). This calculation leads to the conclusion that Piezo1 in an unperturbed RBC should change its mean dome radius of curvature by less than approximately 2 nm as a function of position on the RBC surface.

### Single particle tracking and diffusion analysis

Washed RBCs were adhered to poly-lysine coated 35 mm glass-bottom dishes (MatTeK) and blocked in blocking buffer before incubation with 1:500 dilution of rabbit anti-HA primary (3725; Cell Signaling Technology) for 1 h at room temperature. Cells were then washed in blocking buffer and incubated with 1:20 diluted 40 nm gold-conjugated goat anti-rabbit secondary antibody (Jackson Laboratories) for 1 h at room temperature. Cells were washed in blocking buffer and then DPBS before imaging. The movement of colloidal gold particles was observed at room temperature using differential interference contrast microscopy on an eclipse Ti2 inverted microscope (Nikon) using a 40X/0.65 objective lens and coupled to an Orca Fusion CMOS camera (Hamamatsu). Video sequences were recorded on Nikon NIS-Elements AR5.4.1 software.

Detection and tracking were carried out in Image J imaging analysis software using the TrackMate plugin (Tinevez et al. 2017; Ershov et al. 2021). First, trainable WEKA segmentation (Arganda-Carreras et al. 2017) was employed to generate a model of gold nanoparticles that could be used for spot detection. Detection was then carried out in Trackmate using the Linear Assignment Problem (LAP) tracker. Tracks were exported as XYT coordinates and further analyzed in MATLAB. The MATLAB class msdanalyzer (Tarantino et al. 2014) was used for mean squared displacement analysis of trajectories and estimation of diffusion coefficients.

### Spatial proximity analysis by optimal transport colocalization

Co-immunostained STED images were analyzed using an open-source R code package (Tameling et al. 2021; Tameling and Naas 2021) based on OTC. Sections of 64 x 64 pixels (800 x 800 nm) were picked only if they contained fluorescent signal and were analyzed in parallel to get 95% confidence bands.

## Acknowledgements

Super resolution microscopy work was performed in the Bio-Imaging Resource Center at Rockefeller University, RRID:SCR_017791. The OMX 3D-SIM system was funded by Award Number S10RR031855 from the National Center for Research Resources. This work was also supported at USC by NSF Grant No. DMR-2051681 and by NSF Grant No. DMR-1554716 (to CAH) and at Rockefeller University by NIH grant GM43949 (to RM). R.M. is an investigator of the Howard Hughes Medical Institute.

## Author Contributions

GV and RM conceived the project. GV, PB and AJN designed imaging experiments and GV performed all experiments and analyses. CAH and RM developed calculations of probability distribution of Piezo1 along the red blood cell surface and GV and RM wrote the paper.

## Supplementary figures

**Supplementary Figure 1.**
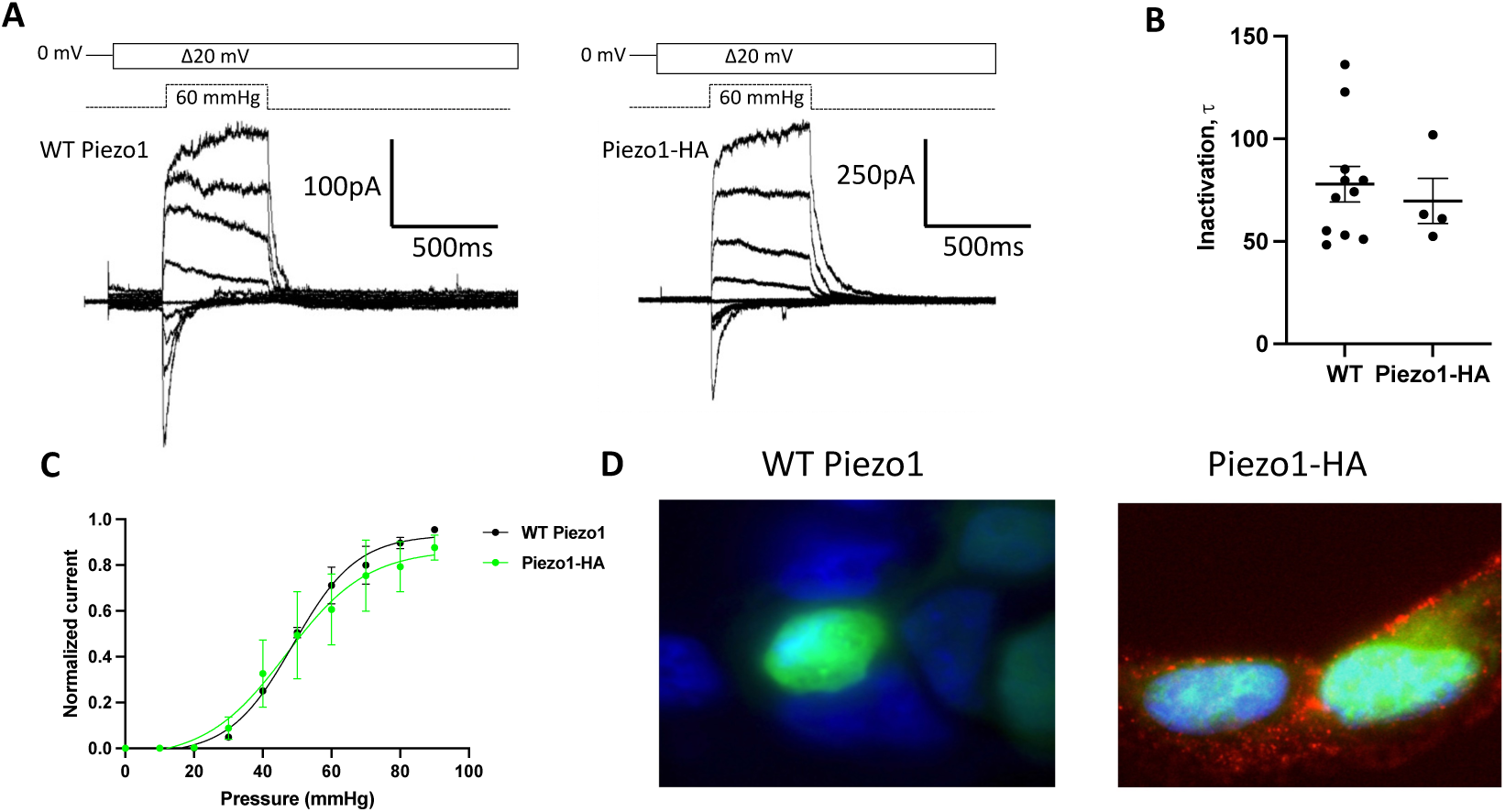
In HEK293 cells HA-tagged Piezo1 behaves like wildtype Piezo1 in electrophysiology recordings and can be specifically detected by immunostaining. **A** Representative traces of currents elicited at 60 mmHg pressure and increasing voltages (in 20 mV steps from -80 to +80 mV) in symmetrical NaCl from excised outside-out patches of HEK293 Piezo1 KO cells overexpressing wild type (left) or HA-tagged mPiezo1 (right). **B** Inactivation time-constants for wild type and HA-tagged mouse Piezo1 with a holding voltage -80 mV and pressure pulses of +60 mmHg. **C** Outside-out patches were held at +60 mV and subjected to 500ms pulses of increasing pressure. Normalized mean current-pressure relations are shown. For each individual patch, currents were normalized to the peak current for that patch. **D** Immunofluorescence in HEK293 Piezo KO cells expressing wildtype or HA-tagged mPiezo1 in a Piezo1-IRES-GFP (green) vector stained with anti-HA primary antibody and Alexa 568-conjugated secondary antibody (red) . Membranes were not permeabilized and DAPI (blue) was used for nuclear staining.

**Supplementary Figure 2.**
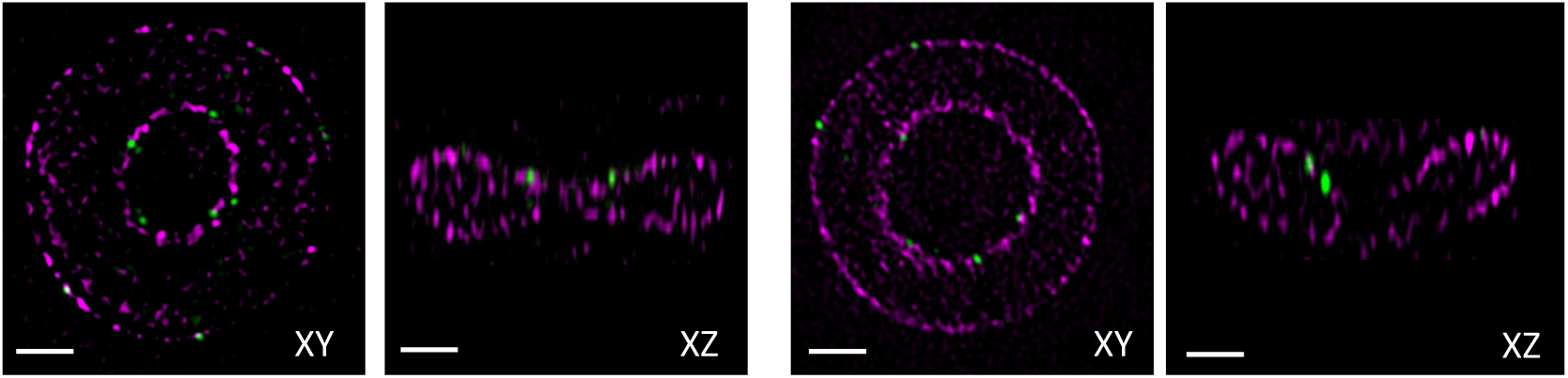
Representative images of XY and XZ two dimensional slice views from SIM images of immunostained RBCs from Piezo1-HA mice. Piezo1 is in green and actin stained by phalloidin is in magenta. Piezo1 spots are observed adjacent to the actin signal, which is a proxy for the membrane in RBCs. Scale bars, 1 μm.

**Supplementary Figure 3.**
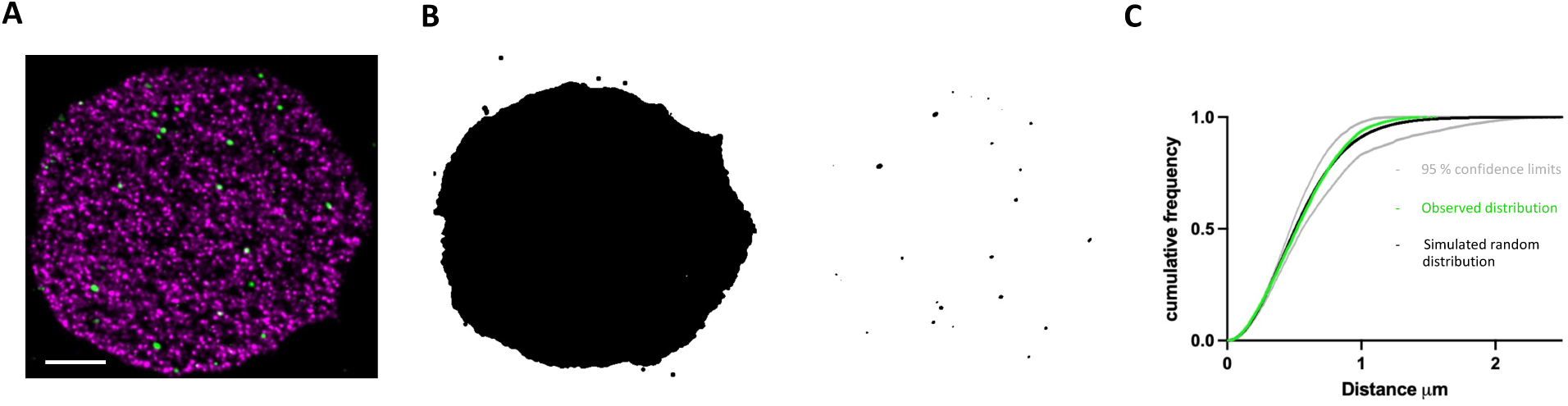
Cluster analysis of fluorescent Piezo1 spots in a 2D STED image of a labelled unroofed RBC membrane. **A** Representative 2D STED image of an unroofed RBC membrane immunostained with an anti-HA tag antibody for Piezo1 (green) and labelled with phalloidin STAR 580 for F-actin (magnta). Scal bar = 1 μm. **B** Binarized mask of the RBC membrane using thresholding of the rhodamine-phalloidin (left) and binarized thresholding of fluorescent Piezo1 spots (right). **C** Cluster analysis using the spatial statistics 2D/3D ImageJ plugin. The cumulative distribution frequency of the distance between a typical position within the masked reference structure and its nearest point in the pattern is plotted on the y axis against the distance, x axis, for a simulated random spatial distribution (black) with 95% confidence limits (grey) compared to the observed experimental distribution (green). There is no statistical difference between the simulated randomly distributed data and observed data.

**Supplementary Figure 4.**
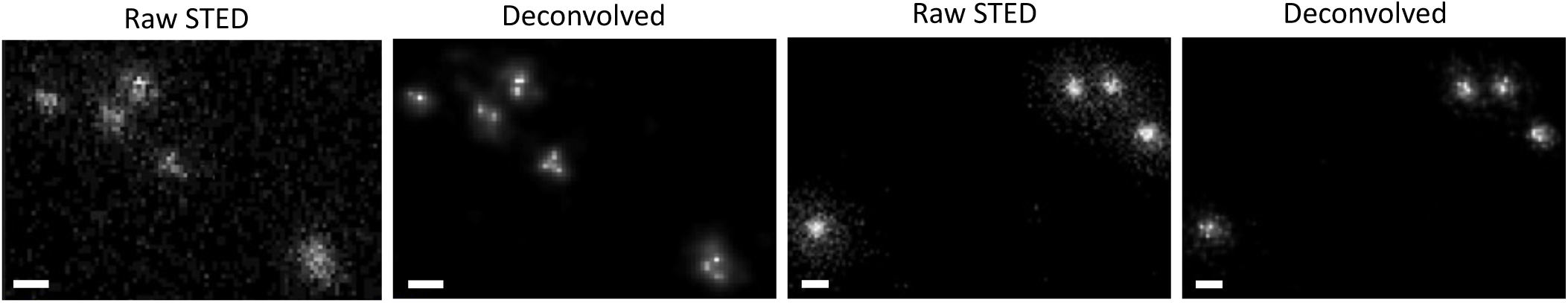
Two representative sets of images of Piezo1 fluorescent spots after STED data collection (left panels) followed by Huygens deconvolution (right panels). Piezo1 spots resolve into singlets, doublets, or triplets, with the bright pixels separated by approximately 25nm. Scale bar, 50 nm.

**Supplementary Figure 5.**
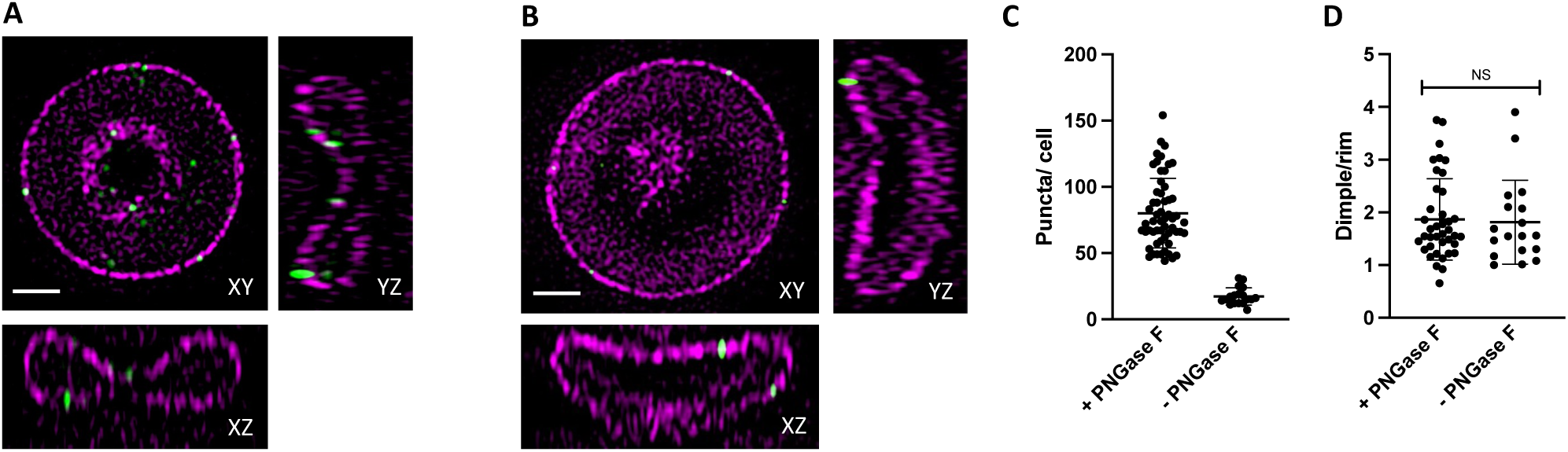
Examples of two cells with different biconcavities and analysis of PNGase F treatment on Piezo1 distribution. **A** 2D slices of a RBC with a biconcavity (measured as the ratio of maximum to minimum heights of the RBC along a central xz slice) of 6.25 and a relative dimple to rim Piezo1 spot density of 2.83. **B** As in *A* but of an RBC with a biconcavity of 1.1 and a relative dimple to rim density of 1.05. Scale bar, 1 μm. **C** Comparison of the number of Piezo1 spots per cell from 3D SIM maximum projections of RBCs prepared with or without PNGase F treatment. For cells treated with PNGase F mean ± SD = 80.2 ± 26.0 spots per cell (N=57 cells) and for cells not treated with PNGase F mean ± SD = 17.3 ± 6.3.0 spots per cell (n=19 cells). **D** Comparison of the Piezo1 spot density in the dimple over rim region of RBCs prepared with or without PNGase F treatment. For cells treated with PNGase F mean ± SD = 1.87 ± 0.76 (N=39 cells) and for cells not treated with PNGase F mean ± SD = 1.81 ± 0.77 (N=18 cells).

**Supplementary Figure 6.**
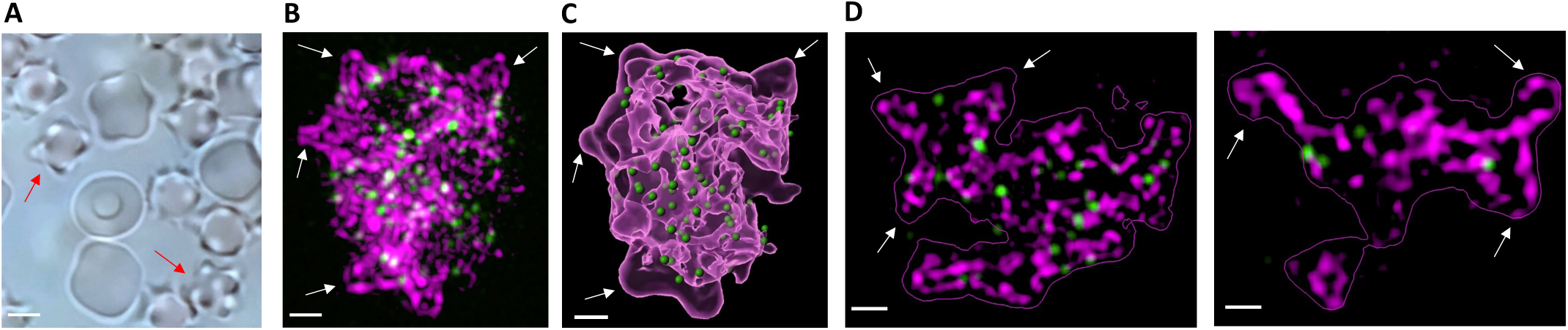
Piezo1 is excluded from membrane regions of high positive curvature in echinocytes. **A** Brightfield imaging of RBCs treated with 10 mM NaSalicylate to generate echinocytes. Two examples of echinocytes are indicated with red arrows. Scale bar, 2 μm **B** Example of an echinocyte labelled with an anti-HA antibody against Piezo1 (green) and rhodamine-phalloidin to label F-actin (magenta). White arrows indicate membrane protrusions of positive curvature where Piezo1 is clearly excluded. Scale bar, 500 nm **C** as in *B* but with the F-actin signal shown as a translucent surface representation (magenta) and the fluorescent Piezo1 spots in green for easier visualization of exclusion of Piezo1 from membrane protrusions. Scale bar, 500 nm **D** left and right are two different Z-slice images from the echinocyte in *B* to illustrate in 2D the exclusion of Piezo1 from membrane protrusions. Scale bar, 500 nm.

**Supplementary Figure 7.**
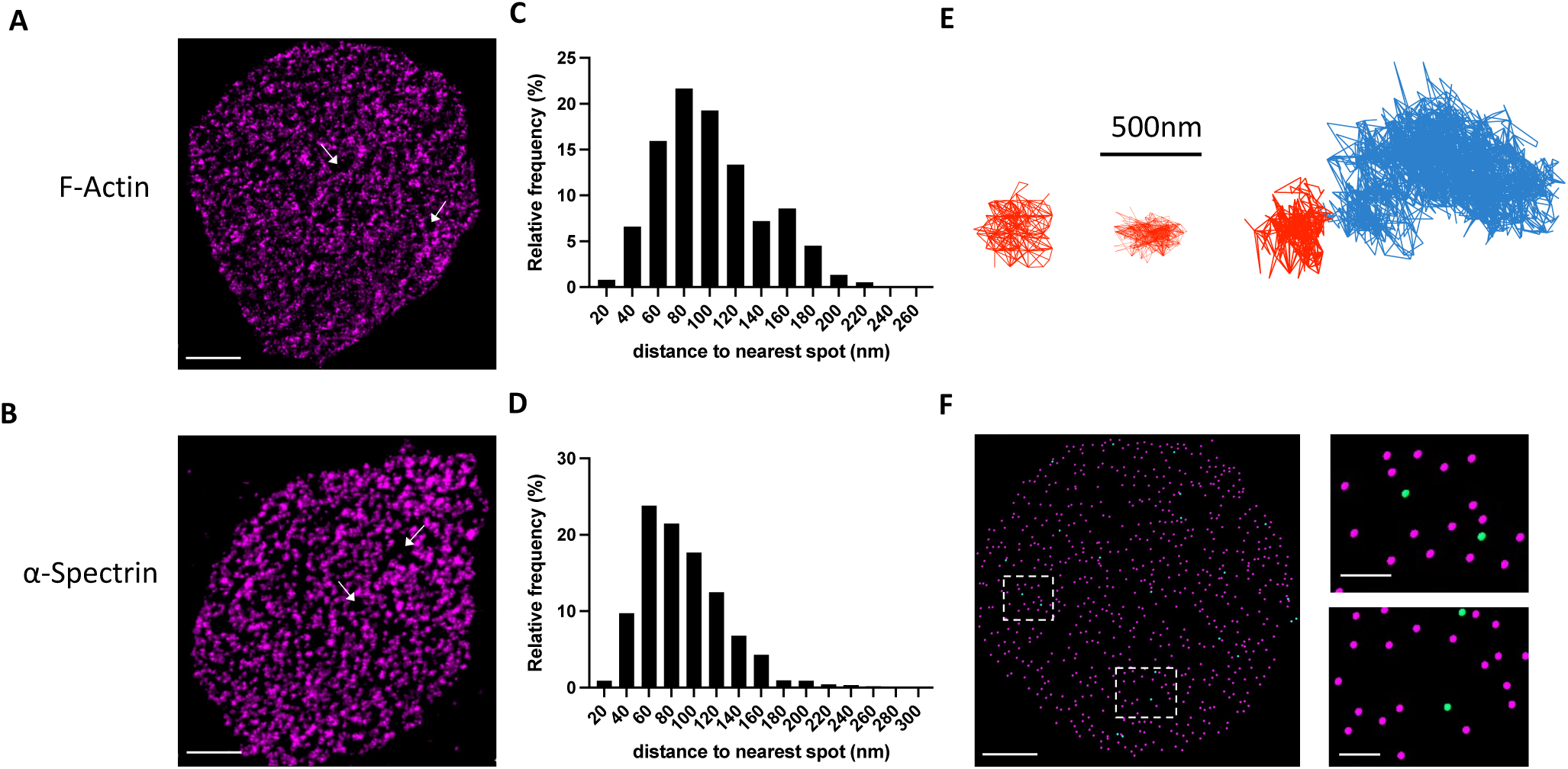
The observed diffusion behavior of Piezo1 is consistent with the organization of the actin-spectrin skeleton in RBCs. **A** 2D STED image of an unroofed RBC stained with phalloidin to label F-actin filaments. White arrows indicate voids deficient in phalloidin staining. These voids are approximately 200- 500nm in size. **B** As in *A* but with immunostaining against α-Spectrin. Scale bar, 1 μm. **C** and **D** Frequency distributions for the nearest-neighbor distance for F-actin (*C*) and α-Spectrin (*D*). **E** Example tracks of Piezo1 exhibiting confined diffusion in spaces < 500 nm (red tracks). The right panel shows a track of a Piezo1 channel that becomes trapped in a < 500 nm confinement radius (red portion of the track) towards the end of the recording. **F** (left) Imaris spot detection was used to detect spots of actin (magenta) and Piezo1 (green) from a 2D STED image of an unroofed RBC. Dashed squares indicate regions of focus (right) where Piezo1 is found in smaller (above) and larger (below) confinement spaces. Scale bars, 1 μm (left) and 200 nm (right).

